# Rescuing Low Frequency Variants within Intra-Host Viral Populations directly from Oxford Nanopore sequencing data

**DOI:** 10.1101/2021.09.03.458038

**Authors:** Yunxi Liu, Joshua Kearney, Medhat Mahmoud, Bryce Kille, Fritz J. Sedlazeck, Todd J. Treangen

## Abstract

Infectious disease monitoring on Oxford Nanopore Technologies (ONT) platforms offers rapid turnaround times and low cost, exemplified by well over a half of million ONT SARS-COV-2 datasets. Tracking low frequency intra-host variants has provided important insights with respect to elucidating within host viral population dynamics and transmission. However, given the higher error rate of ONT, accurate identification of intra-host variants with low allele frequencies remains an open challenge with no viable solutions available. In response to this need, we present Variabel, a novel approach and first method designed for rescuing low frequency intra-host variants from ONT data alone. We evaluated Variabel on both within patient and across patient paired Illumina and ONT datasets; our results show that Variabel can accurately identify low frequency variants below 0.5 allele frequency, outperforming existing state-of-the-art ONT variant callers for this task. Variabel is open-source and available for download at: www.gitlab.com/treangenlab/variabel.

## Main Text

Oxford Nanopore Technology (ONT) has become a dominant technology for rapid sequencing of COVID-19 patients due to its low cost and relatively simple preparation methods^1^. ONT datasets have proliferated during the pandemic; there are now well over 100,000 sequenced COVID-19 samples from ONT alone in the NCBI SRA database and over a half a million SARS-CoV-2 genomes assembled from ONT^2^. Intra-host variation of COVID-19 reveals important information about many aspects of the disease, such as future variants of concern and the response to different treatments^3,4^. When SARS-CoV-2 infects a human host, a combination of viral and host proteins facilitates the replication of the virus ^5^. Intra-host variants then arise during the expansion of the intra-host viral population and homologous recombination ^6^, some of which may be biologically relevant ^7^. Given the ubiquity of COVID-19 ONT data, the goal of our study is to explore the use of widespread ONT data for detection of intra-host variation to elucidate currently “hidden” biology. However, due to the relatively high error rate of ONT, ranging from 5%-15%, true variation within hosts is obscured by sequencing errors contained in the raw data ^8^. Our assumption is that the allele frequency of true SNV within a sample is subject to change across samples, while those of sequencing errors are independent of the sample and thus are highly stable/similar. This is especially the case for similar bascaller and flow cells versions. Multiple studies have shown that the allele frequency of a true variant would experience a significant change over time or over samples collected from different patients ^9–12^. Indeed, the size of the population of SARS-CoV-2 virions within a host undergoes exponential growth post infection, increasing from a handful of virions to one billion virions or more ^13^. Furthermore, the vast majority of sequencing errors in ONT data are deletions, related to homopolymer regions where the same nucleotide occurs consecutively, or low-complexity regions ^14–16^.

Current read-based error correction and polishing methods for ONT data primarily target genome assembly ^17,18^. Raw read polishing has proven to be extremely effective in generating high quality assemblies, however, information supporting low frequency (less than 0.5) intra-host variants is almost always lost during the process. An alternative approach to preserving intra-host variation during error correction involves integrating haplotype information into the assembly step^19, 20^. Strainline uses a combination of local De Bruijn graph assembly and overlap extending to generate haplotype genomes^19^. CliqueSNV constructs haplotype sequences by recognizing linked SNVs that are supported by a single read^21^. While both methods claim to assemble genomes at strain level resolution, haplotype phasing from ONT sequencing protocols for SARS-CoV-2 is challenging due to the limited read length from amplicon sequencing (250bp-500bp) ^22^, uneven coverage, and susceptibility to bias from single nucleotide variation. Furthermore, sequencing error in ONT data is context dependent ^16,23^.

Here, we present Variabel, a novel variant call filtering tool that is able to recover intra-host variants from ONT data alone, for the first time, by exploiting the tendency of true variants to change in allele frequency across samples. The key concept behind Variabel is that by leveraging information from viral population dynamics, we can distinguish the true variants from sequencing errors caused by ONT by comparing samples collected across different time points or samples collected from different patients. Variabel is constructed as a series of filters that operate on the variant call format (VCF) files returned by an existing variant caller. **Figure 1** illustrates the variant calling workflow and the algorithms of Variabel. It includes an allele frequency variation filter, which identifies true variants that are shared across different samples (see **Figure 1B**) and an insertion/deletion (indel) filter that identifies false indel calls based on Shannon’s entropy values of the region near indel sites (see **Figure 1C**).

**Figure 1.**
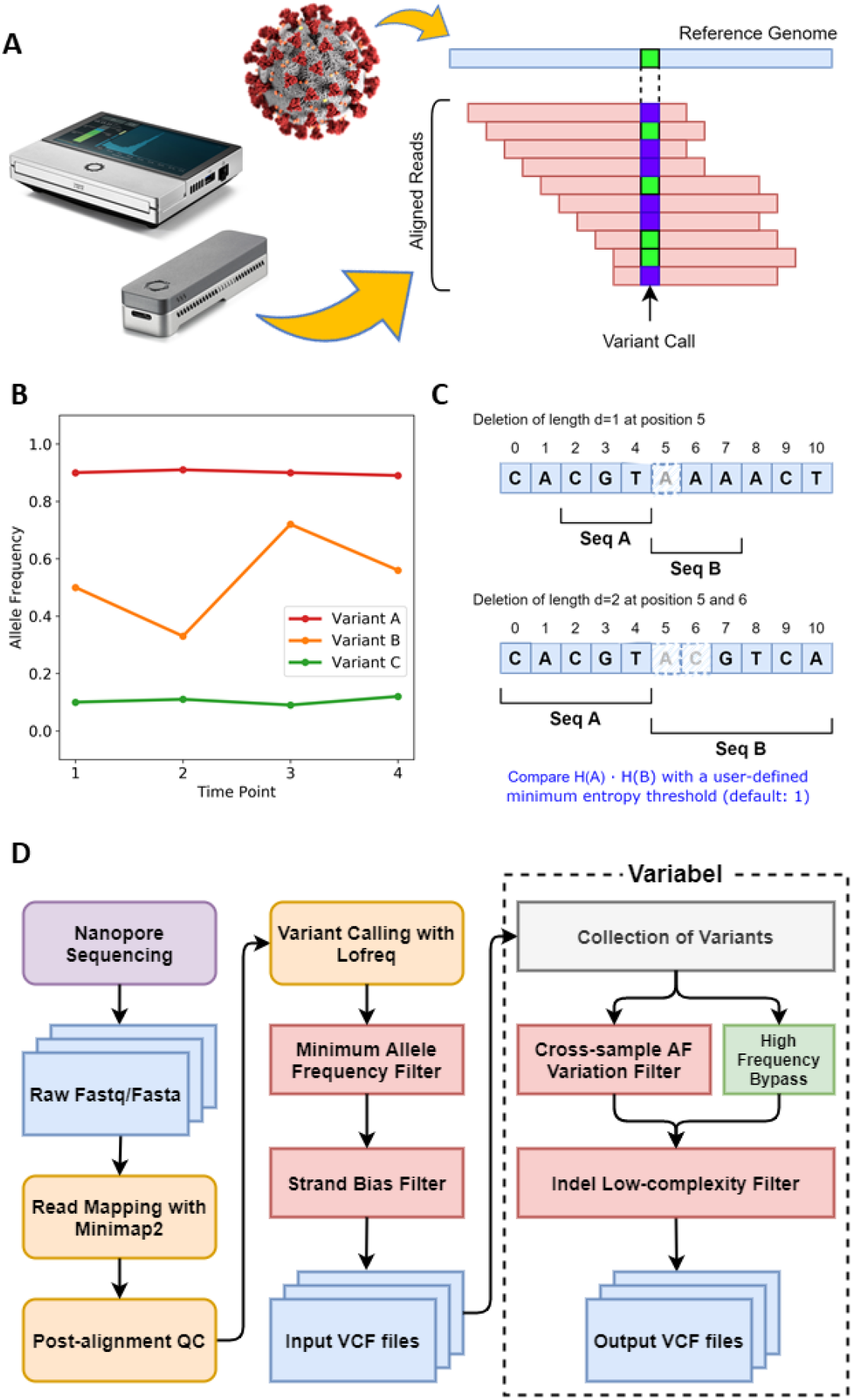
Illustration of Variabel algorithm and workflow. **a)** Sequencing reads from ONT are aligned to the reference genome of SARS-CoV-2 with Minimap2, then variants are called based on the alignments using Lofreq. **b)** Cross-sample AF variantion filter identifies variants that are shared between samples. Variant calls with maximum AF less 0.65 and maximum AF variation less than 0.05 are classified as false calls. In this example, variant A and B pass the filter while variant C fails. **c)** Low-entropy filter calculates the Shannon’s entropy H for subsequences (Seq A and B) of the reference genome around the position where the indel call occurs. The length of the subsequences is determined by the length of the indel. Product of two entropy of the subsequences is used to determine whether the indel is a false positive or not. **d)** Workflow of Variabel for detecting intra-host variants for ONT sequences.

We evaluated Variabel via two ONT datasets: 1) a time-series COVID-19 dataset and 2) a cross-patient dataset ^4,24^. Importantly, samples in both datasets are from studies sequenced with both ONT and Illumina. The time-series dataset contains samples taken from an immunocompromised COVID-19 patient over the course of three months^4^, where 18 pairs of Illumina and ONT sequencing runs passed quality control. The cross-patient dataset includes 154 SARS-CoV-2 positive samples collected from patients, and 103 pairs of Illumina and ONT sequencing runs passed our quality control. We selected these two experimental datasets for evaluation as: i) the time-series dataset allows us to track individual changes in allele frequencies over time for a specific patient, and ii) the cross-patient datasets allow us to explore the utility of Variabel on more readily available SARS-CoV-2 datasets. Variant calls on Illumina sequencing runs by Lofreq ^25^ are used as a benchmark in our calculation of precision,recall, and f-score. In the benchmark, the time series dataset contains 865 substitutions and 133 indels, and the cross patient dataset has 3,236 substitution and 990 indels. We also ran Clair3 on the same ONT sequencing runs for benchmarking purposes. While Clair3 is not explicitly designed for virus SNV calling, it represents a state-of-the-art ONT variant caller ^26^.

Illumina sequencing produces highly accurate reads, which are ideal for intra-host variant calling. On the other hand, while variant calling on ONT sequencing data offers faster turnaround time and is not limited to sequencing centers, it is much more challenging due to a higher error rate in both the sequencing and base calling process. Most of the previously reported intrahost variants have allele frequencies of >0.02 and less than 0.15, which is well above the Illumina error rate but *exactly within* the expected ONT error rate, highlighting the dichotomy of using one or the other for identification of low frequency intra-host variation. Our results highlight that Variabel is able to call variants in ONT data with high precision. **Figure 2** indicates the positions and minimum allele frequencies of variants called by Lofreq and Variabel. By comparing the variant calling results before and after applying Variabel on both time series dataset (**Figure 2A**) and cross patient dataset (**Figure 2B**), we found that Variabel is able to remove the majority of the false positive calls caused by the sequencing errors of ONT data. The number of variants that are exclusively found in ONT data (marked in green) drop dramatically, while most of the true variants which are found in both Illumina and ONT reads passed the filters.

**Figure 2.**
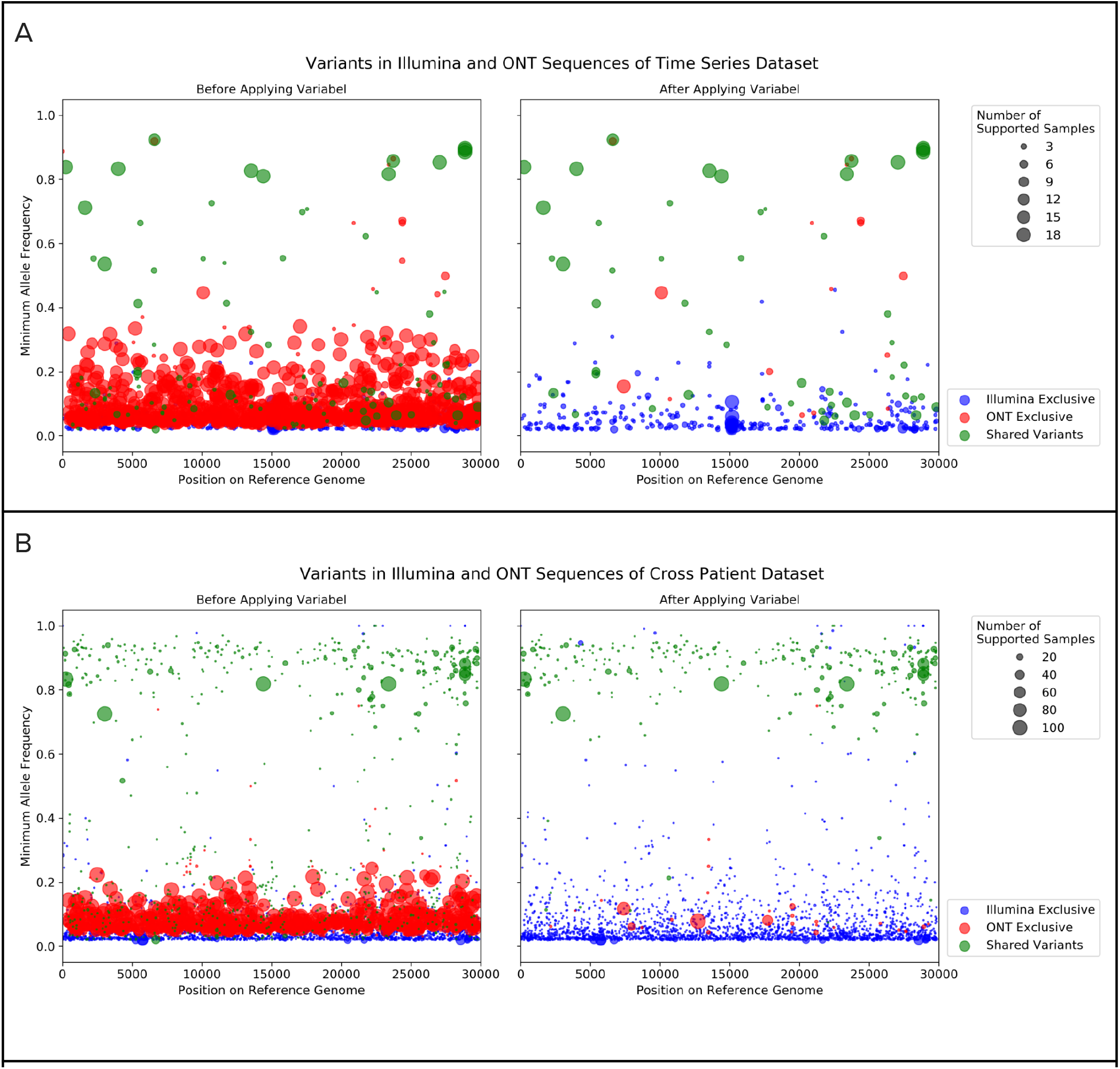
Variants called by lofreq with Illumina and ONT sequences before and after applying Variabel. In each of the subfigures, the left plot shows the variant calls before applying Variabel and the right plot shows the variant calls after applying Variabel. The x-axis shows the position of the variant on the reference genome. The y-axis shows the minimum allele frequency of the variant found in multiple samples. Variants found in the Illumina sequences only are marked in blue, and variants found in the ONT sequences only are marked in red. Variants that are shared between both Illumina and ONT data are shown in green. The size of the dot represents the number of samples supporting the variant. **a)** Variant calls of the time series dataset. **b)** Variant calls of the cross patient dataset.

We also benchmarked Variabel with Clair3. **Figure 3A** shows a Venn diagram of variant calls from Variabel and Clair3 compared to Illumina variant calls for 18 time series samples. Out of 474 variant calls made by Variabel, 405 (85.44%) of them are considered true positive calls since they are also found in the Illumina data. Clair3 had a lower number (388) of true positive calls compared to Variabel, and Clair3 had 412 false positives while Variabel only had 69.

**Figure 3.**
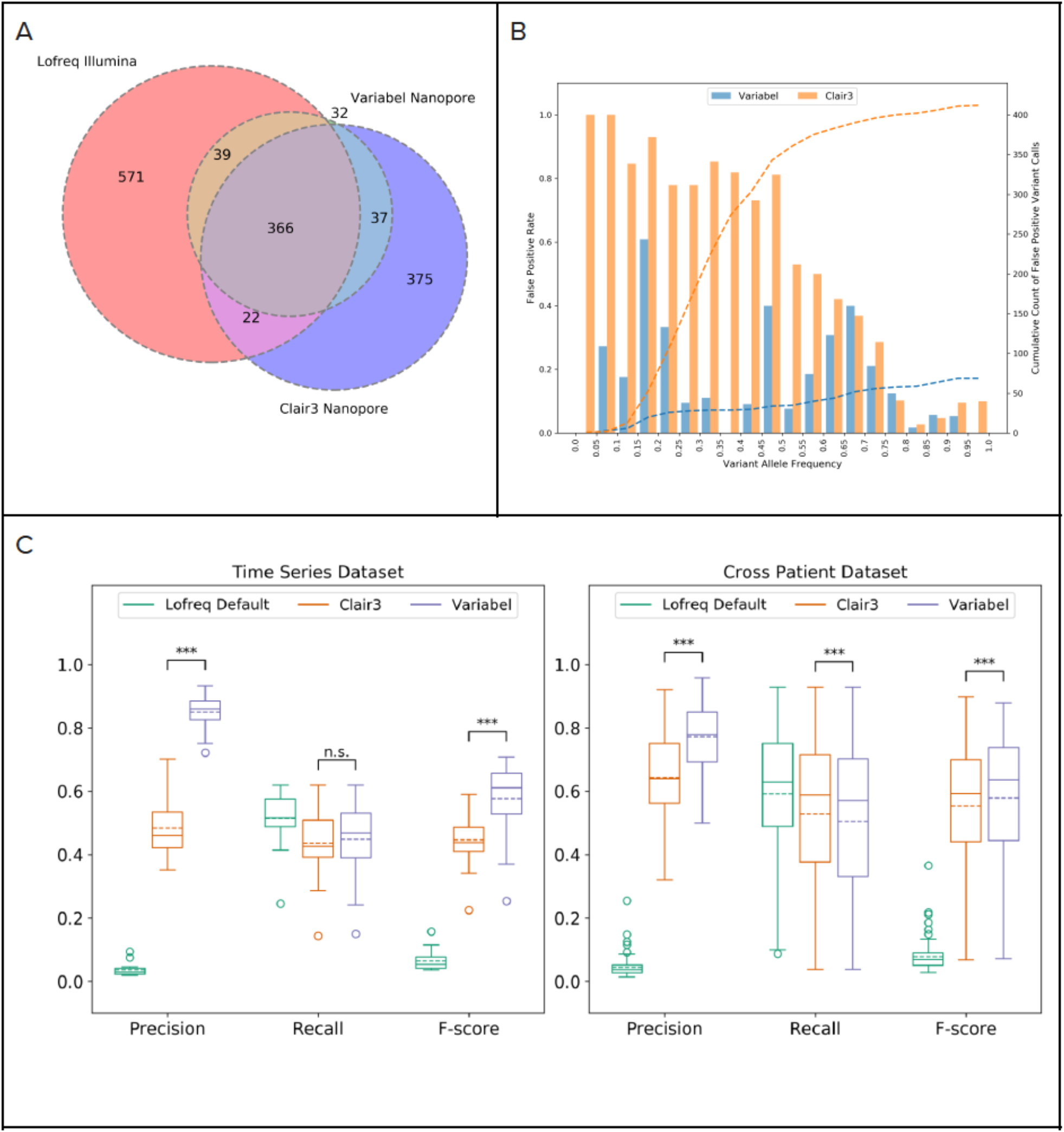
Intra-host variant detection on time series dataset. **a)** Venn diagram showing counts of variant calls shared between Lofreq on illumina sequencing runs and Variabel and Clair3 on nanopore sequencing runs on the same samples from time series dataset. b) False positive rates at different variant allele frequencies and cumulative count of false positive variant calls of Variabel and Clair3 for time series dataset. c) Precision, recall, and f-score comparison of Lofreq default, Clair3, and Variabel on both time series dataset and cross patient dataset. Significance between Clair3 and Variabel were calculated using the two-sided paired t-test. Significance labeling: n.s.(P>0.05), *(P0.05), **(P0.01), ***(P0.001).

Importantly, Variabel is able to rescue true variants in the low frequency domain. **Figure 3B** shows the false positive (FP) rates at different variant allele frequencies and cumulative density of FP variant calls from Variabel and Clair3 for time series dataset. First, we see that for ultra-low frequency variants (less than 0.1 allele frequency), Clair3 has a FP rate of 100% (all variants identified are false positives), while Variabel’s FP rate for these ultra-low frequency variants is less than 0.2. Next, we observed that Variabel has much lower FP rate on average for variants with allele frequency below 0.65 compared to Clair3. The peak FP rate for Variabel occurs at allele frequency between 0.15 and 0.2, which is associated with Nanopore sequencing errors. Cumulative count plot of FP variant calls shows that Variabel has a close to uniform distribution of false calls along different allele frequencies. On the other hand, more than 80% of the FP calls from Clair3 have allele frequencies less than 0.5.

**Figure 3C** shows precision, recall and f-score for variant calls generated by different methods on both time series and cross patient datasets; shown are the LoFreq default output with 0.02 minimum allele frequency filter, Variabel, and Clair3. For all 18 samples that passed quality control from the time-series dataset, applying Variabel resulted in a significant mean precision increase from 0.036 to 0.850, and a mean f-score increase from 0.065 to 0.578. When applied to the same data, Clair3 had a mean precision of 0.483 and a recall of 0.446, noticeably underperforming Variabel. Both Variabel and Clair3 had similar mean recall (0.448 for Variabel and 0.436 for Clair3). Similar results can also be found in the cross-patient dataset: for all 103 samples that passed quality control (see methods), the mean precision increased from 0.044 to 0.772 after applying Variable, which is significantly higher than the Clair3 mean precision of 0.641. The mean f-score is 0.578 for Variabel and 0.553 for Clair3. **Figure 4A** shows the Venn diagram of variant calls for 103 cross patient samples: although Vairabel had 356 false positive calls on this dataset, Clair3 had twice as many (753 total).

**Figure 4.**
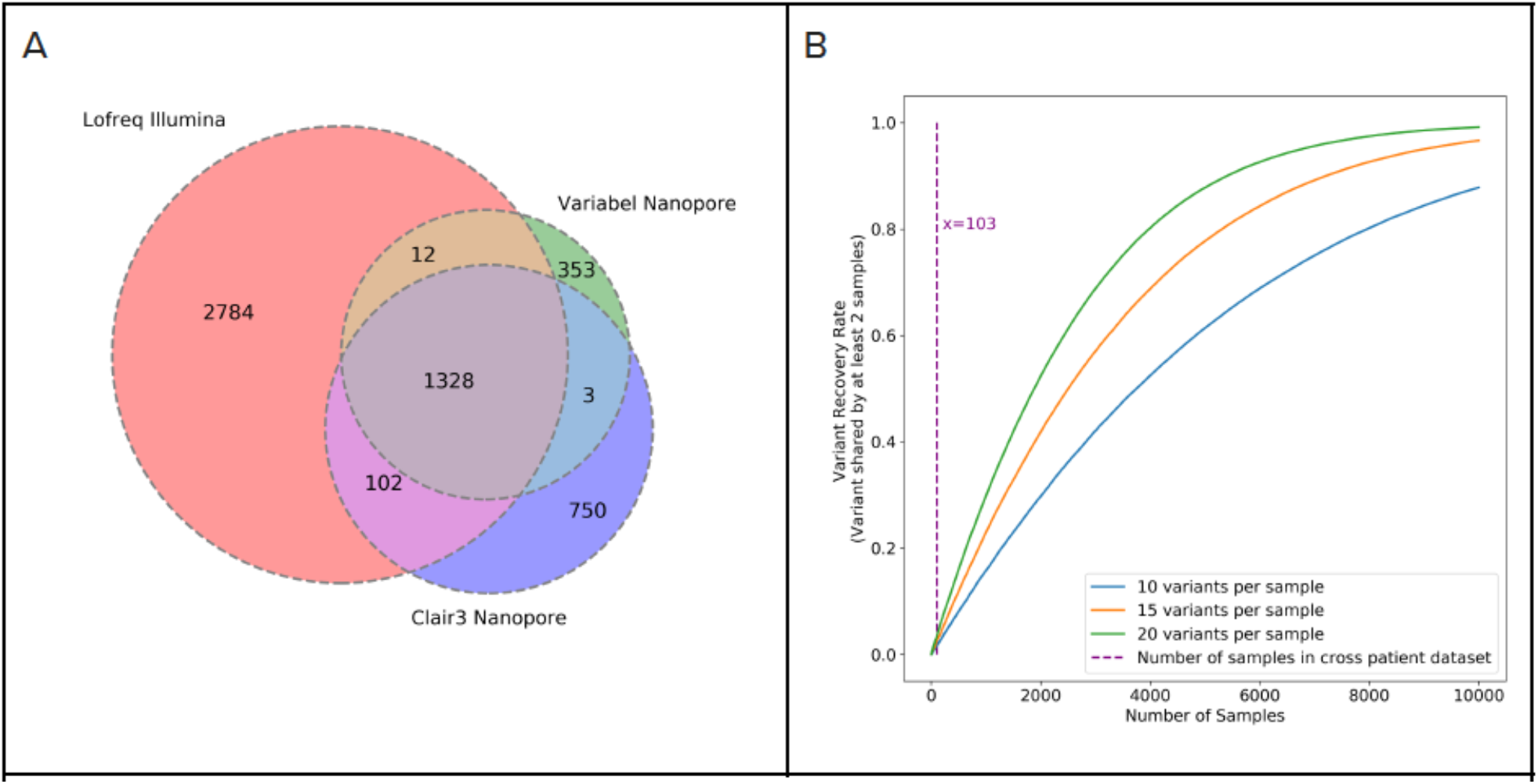
Intra-host variant detection on cross patient dataset. **a)** Venn diagram showing counts of variant calls shared between Lofreq on illumina sequencing runs and Variabel and Clair3 on nanopore sequencing runs on the same samples from cross patient dataset. **b)** Simulation of fraction of shared variants recovered from different sizes of collections of SARS-CoV-2 samples.

Our results highlight that it is possible to accurately identify emerging intra-host variations using ONT sequencing alone. This enables a fast and accurate variant prevalence utilizing the scalability and turnaround time of ONT that is already in place around the world. Variabel uses the variant frequency information and entropy filtering to distinguish true intra-host variants from ONT sequencing error. This is well maintained in the time-series data. Furthermore, our experiments have shown that the usage of Variabel can be extended to cross patient datasets, which strongly hints at broader applicability of our approach to the vast amount of ONT based COVID studies.

One of the main limitations of Variabel for cross-patient studies is that the same variant must be observed in at least two samples to activate allele frequency variation filtering. As expected, we observed a drop in average precision in the cross patient dataset compared to time-series dataset, since samples collected from different patients are less likely to contain shared variants. Rescuing low frequency intra-host variants is far more challenging for cross patient data compared to longitudinal data, and the distribution of allele frequencies of true positive variants found by Variabel in the cross-patient dataset clustered above 0.6 while allele frequencies of true positive variants spans in much wider range in the time series dataset (see **Figure 2**). Based on a simple simulation (see **Figure 4B**) we calculate that approximately 10,000 samples would be required to recover most of the intra-host variants if we assume variants occur randomly along the genome of SARS-CoV-2. Similarly, we also expect a small drop in performance of Variabel if the time series data included fewer samples (e.g., 2-5). Both scenarios could be improved by leveraging a centralized data depository of low frequency SNV for SARS-Cov-2. Follow-up studies can then leverage this resource to assess and evaluate the biological importance of observed low frequency variants within and across hosts over time. However, established COVID databases such as GISAID are limited only to consensus level sequences ^27,28^, which might be a limiting factor going forward in this or future pandemic or outbreaks.

In conclusion, Variabel is the first method explicitly designed to identify low frequency intra-host variants directly from ONT data in viral populations. This represents both an important advance for the field and will facilitate the tracking of intra-host variation in COVID-19 positive patients.

## Methods

### Dataset descriptions

Two datasets were used to validate the performance of Variabel. The time series dataset is a longitudinal dataset containing respiratory samples collected from a single SARS-CoV-2 positive patient with immunodeficiency at 23 different time points across a 101-day period ^**4**^. All samples were sequenced with MinlON (Oxford Nanopore Technologies). Among those samples. 20 samples were deep sequenced using the lllumina platform. The ONT data downloaded from NCBI SRA database showed that the quality scores of the reads are corrupted. All bases were assigned with the same quality score”?”. The cross-patient dataset contains 154 SARS-CoV-2 positive samples that are collected from different patients and sequenced using both illumina and nanopore platforms ^**24**.^ All raw sequencing runs from both datasets are publically available on NCBI SRA database.

### Quality control and read alignment

We performed pre-alignment quality control on all illumina sequencing runs using fastp (v0.20.1). ^**29**^ with the following command. fastp --detect_adapter_for_pe --cut_front, --cut_window_size 4 --cut_mean_quality 15 --length_required 15 --qualified_quality_phred 15 --unqualified_percent_limit 40 --n_base_limit 5 --low_complexity_filter. Read alignment for illumina sequences was done using bwa mem (v0.7.17-r1188) paired end mode with default parameters. ^30^ For nanopore Read alignment for nanopore sequences was done using minimap2 (2.20-r1061) with preset map-ont.^**31**^ Alignment files are sorted using samtools (v1.11) ^32^. We also performed post alignment quality control by calculating breadth and depth of genome coverages with samtools depth. For each pair of illumina and nanopore sequencing runs which were generated from the same sample, both sequencing runs must have breadth of genome coverage no less than 0.9 and mean depth of coverage no less than 500, otherwise both sequencing runs are excluded from our experiments.

### Variant Calling

We used lofreq (v2.1.4) to call variants for illumina samples. This is done by first inserting indel quality score into the BAM files using command “lofreq indelqual --dindel”, and then call variants including insertions or deletions (indels) with command “lofreq call --no-default-filter --call-indels”. At last, we applied the strain bias filter and removed any variants with allele frequency below 2% with command “lofreq filter --cov-min 2 --af-min 0.02 --sb-alpha 0.01 --sb-incl-indels”.^25^. Both Variabel and Clair3 were used to call variants from nanopore data. We used Clair version 3 [https://github.com/HKU-BAL/Clair3] ^26^ to identify SNVs and indels in samples using default parameters. We used the training data set specified for ONT and set the --chunk_size to 29,903.

To call variants with variabel, we first stripped the quality score from the nanopore data in order to force Lofreq run with its EM algorithm. Then we insert the indel quality score into the BAM files using command “lofreq indelqual --uniform 16”. After that, we used the same command as processing illumina data to call variants and to filter variants with great strain bias or with allele frequency below 2%. The collection of VCF files is used as input for Variabel. First, Variabel performs the cross-sample allele frequency variation filtering. It examines each one of the VCF files, identifies variants that are shared between samples, and records their allele frequencies. Any variant with maximum allele frequency less than 0.65 and maximum variation of 0.05 or less across all the samples in which the variant existed is classified as false calls and is eliminated. Variabel then applies a low-entropy filter to any indels that occur in regions of low-complexity. This is designed to eliminate nanopore homopolymer errors that primarily occur in areas with short repeats. Assume a deletion is called at position i on reference genome s with length d, the filter checks the product of the Shannon entropy of the substring s[i-2d: i+1] and the Shannon entropy of the substring s[i+1: i+1+3d]. If the value of the product is less than the user defined threshold (default: 1), the variant is classified as false calls and is eliminated. The variants that pass both cross-sample allele frequency variation filter and low-entropy filter are collected and output in VCF format.

### Validation

Since all samples we included in our analysis have both illumina and nanopore sequencing runs in high quality, we used the lofreq calls generated from the illumina data as a ground truth to evaluate precision, recall, and f-score of Variabel and Clair3.

## Authors Contributions

All authors conceived the experiments, analyzed the results, and reviewed the manuscript. YL, BK, JK, MM conducted the experiments. YL and JK wrote the code. FS and TT managed the project.

## Acknowledgements

We would like to thank the contributing authors of the paired Illumina and ONT SARS-CoV-2 sequencing data which was instrumental for highlighting the benefits of Variabel on ONT data. MM and FS were supported by the National Institute of Allergy and Infectious Diseases (Grant#1U19AI144297). TT was supported in part by the National Institute of Allergy and Infectious Diseases (Grant#1P01AI152999-01). TT and YL were supported in part by the C3.ai Digital Transformation Institute COVID-19 award and Centers for Disease Control (CDC) contract 75D30121C11180.

